# Characterisation of a touchscreen-based Go/No Go task for assessing cognitive judgement bias in mice: a new translational tool for affective state disorder drug screening

**DOI:** 10.1101/2025.01.20.633915

**Authors:** L Lopez-Cruz, BU Phillips, LM Saksida, CJ Heath, TJ Bussey

## Abstract

A major obstacle in the pre-clinical study of mood-related disorders and novel affective state therapeutic evaluation is the lack of animal models that fully recapitulate human symptomatology.

In this study, we developed a touchscreen-based Go/No Go cognitive judgement bias (CJB) task for mice. In this task, animals first learned to discriminate between two visual stimuli displayed on the touchscreen: one associated with a reward (S+) and one associated with a time-out and flashing house light (S−). Once mice learned to respond to the S+ and to withhold responding to the S− consistently, a set of four ambiguous stimuli ranging in visual similarity to the S+ and S− stimuli were randomly interspersed in the stimulus presentation sequence. Responses to these ambiguous stimuli were interpreted as a greater expectation of positive (‘optimistic bias’) or negative (‘pessimistic bias’) outcomes as a function of their similarity to the S+ or S− stimuli.

The acute administration of the SSRIs fluoxetine and citalopram and the 5HT-2C receptor antagonist SB204048 did not produce any effects on CJB task performance. However, the noradrenaline/dopamine reuptake inhibitor, bupropion, increased responses to the ambiguous stimuli consistent with induction of an ‘optimistic bias’ and the pro-depressant tetrabenazine yielded the opposite effect.

This study underscores the capacity of mice to respond to visually ambiguous stimuli in an ambiguity-dependent manner, a phenomenon observed across various species, including humans. Furthermore, it establishes and validates an operant behavioural task to assess CJB in mice delivered using the touchscreen platform which has significant intrinsic cross-species translational potential.

## Introduction

Mood disorders are among the most prevalent psychopathologies, with depression the leading contributor to the global burden of disease (Lin et al. 2014). The preclinical study of mood disorders, and specifically of depression, has been challenging historically because of their complexity, the heterogeneity of symptoms between and within patients, and the intricate, subjective nature of human emotions (Planchez et al. 2019). Combined, these factors have contributed to the difficulty in developing animal models that accurately recapitulate these clinical conditions.

Although there are preclinical tools available for measuring and modeling some core mood disorder-related symptomatology such as anhedonia (Scheggi et al. 2018), anergia and lethargy (Salamone and Correa 2018), as well as additional symptoms like changes in weight and appetite, some other pivotal symptoms like sadness or depressed mood are difficult, if not impossible, to address in animal models. Traditional approaches for evaluating depressive-like symptomatology are based on the assessment of despair-like behaviors, as in the forced swim test (FST) and tail suspension test (TST) (Porsolt et al. 1977; Kulkarni and Mehta 1985; Steru et al. 1985; Castagné et al. 2011) which consist of placing animals in unescapable, stressful situations. However, the behavioral endpoints of these assays (i.e., immobility, swimming, or struggling) do not reproduce the clinical condition. Moreover, these methods, although used as gold standards for antidepressant screening, have recently raised both scientific and animal welfare concerns (Commons et al. 2017; Sewell et al. 2021; Trunnell and Carvalho 2021).

To address the issues associated with the preclinical assessment of mood-related symptoms, the measurement of cognition-related processes that are also compromised in depression has been suggested (Pan et al. 2019; Panchal et al. 2019). In particular, the measurement of cognitive judgement bias (CJB), which indexes how specific emotions influence biases in information processing or interpretation, has been proposed as an indirect approach to measuring changes in mood (Paul et al. 2005; Anderson et al. 2012; Hales et al. 2014; Roelofs et al. 2016; Robinson and Roiser 2016). Negative cognitive biases (manifested as interpreting ambiguous situations or stimuli as negative) have been observed in patients with depression and in participants with an increased vulnerability to this psychopathology (Mathews and MacLeod 2005; Hayward et al. 2005), underlining the clinical relevance of this metric. Changes in CJB have also been observed in other mood disorders (MacLeod and Mathews 2012), suggesting potential transdiagnostic utility.

Promisingly, previous studies have demonstrated the existence of CJB across species and assays targeting this construct have been developed in mice (Krakenberg et al. 2019, 2020), rats (Harding et al. 2004; Enkel et al. 2010; Krakenberg et al. 2019, 2020), and non-human primates (Bethell et al. 2012; Bethell and Koyama 2015)). Taken together, these tasks make use of a multitude of designs and reward/punishment contingencies and involve engagement with and discrimination between stimuli varied across a range of domains and parameters (e.g. tones or visual stimuli presented with varying properties and in distinct spatial locations). That it is possible to engage CJB across multiple species via such highly varied paradigms arguably speaks to the robustness of the construct, but this heterogeneity of assay design also makes inter-study and inter-species comparisons challenging. Further, while each assay has been optimized for use in the particular species in which it was characterized, successful translation to humans may also be problematic for a variety of reasons. The use of touchscreen-based devices for behavioural assessment addresses both of these potential issues through rigorous standardization of approach and stimulus parameters, the capacity for the assays to be driven within session by multiple visual stimuli of various sizes and shapes that can be presented dynamically across multiple spatial locations (thereby broadening flexibility of design relative to traditional operant manipulanda-based approaches) and an established record of preclinical to clinical assay translation using these systems (Nithianantharajah and Grant 2013; Nithianantharajah et al. 2015; Heath et al. 2019; Lopez-Cruz et al. 2021; Krahn et al. 2024).

While touchscreen-based CJB tasks for mice have been characterized previously (Krakenberg et al. 2019), in this work we intend to optimize and validate a mouse CJB paradigm the design principles of which have been previously utilized in non-human primates (Bethell et al. 2012). Successful characterization of such a CJB assay in mice would enable evaluation of the capacity of this species to respond across a varying range of ambiguous stimuli in the same behavioural session, as reported previously in other species including humans (Enkel et al. 2010; Bethell et al. 2012; Anderson et al. 2012) while, unlike other touchscreen based-tasks, using a single location for both optimistic and negative responses, using more than one ambiguous stimulus and using a valence-based contingency (reward vs. no reward) instead of reward-based continencies (high reward vs. low reward) as positive and negative discriminators (Krakenberg et al. 2019, 2020).

In the Bethell et al. (2012) non-human primate CJB task, animals learned to discriminate between two visual stimuli (rectangles of different lengths) displayed on the screen. The “positive stimulus” was associated with a reward, and the “negative stimulus” was associated with the absence of reward and a time-out (a 16 seconds blue screen and a 2 second white noise).. Animals had to respond to the positive stimulus (i.e., ‘Go’ response) while withholding responses to the negative stimulus (i.e., ‘No Go’ response). Once animals learned to discriminate, they were presented with intermediate-length rectangles (ambiguous stimuli) which varied in similarity to the positive and negative stimuli.

With respect to validation, Bethell et al. (2012) reported that animals in enriched housing conditions made more responses to the ambiguous stimuli, indicating a greater expectation of positive outcomes (‘optimistic bias’), whereas those subjected to stressful vet checks made less responses to ambiguous stimuli, indicating a higher expectation of negative outcomes (‘pessimistic bias’). Crucially, from a cross-species perspective, this pattern of responses has also been observed in humans with depression in tasks using the same Go/No Go architecture which feature humanized stimuli like words or emotional faces (Erickson et al. 2005; Schulz et al. 2007), suggesting significant translational potential for this approach.

To validate the new Go/No Go CJB task in mice reported here, its sensitivity to changes in mood or affective-like state are evaluated using a range of pharmacological manipulations with established neurochemical profiles including the Selective Serotonin Reuptake Inhibitors (SSRIs) citalopram and fluoxetine (Anderson et al. 2013; Stuart et al. 2013), the noradrenaline and dopamine reuptake inhibitor bupropion (which has been shown to decrease negative bias in humans (Harmer et al. 2009), the 5HT-2C receptor antagonist SB204048 (Phillips et al. 2018) and the vesicular monoamine transporter 2 blocker, tetrabenazine (TBZ), which has been shown to have pro-depressant effects (Stuart et al. 2017; Hales et al. 2022; Carratalá-Ros et al. 2023).

## MATERIALS AND METHODS

### Animals

Adult male C57BL/6 mice (n=45) (Charles River Laboratories, Margate, UK), 8-10 weeks old at the beginning of the experiments, were housed in groups of 4 (lights off 07:00, lights on 19:00). After facility habituation for 7 days, all animals were weighed for 3 consecutive days to establish mean free-feeding weights. Mild food restriction was initiated and sustained at 85-90% of free-feeding weight by daily provision of specific amounts of standard laboratory chow (RM3, Special Diet Services). Drinking water was available ad libitum throughout the study. All animals were tested/trained once daily 5-7 days a week during the dark phase. All procedures were performed in accordance the United Kingdom Animals (Scientific Procedures) Act (1986) and the United Kingdom Animals (Scientific Procedures) Act (1986) Amendment Regulations 2012 and had been reviewed and approved by the University of Cambridge AWERB.

### Drugs

Bupropion (Tocris Bioscience, Bristol, UK) was dissolved in 0.9% (w/v) saline at the required concentrations and administered intraperitoneally (IP) with a 30-min delay between injection and the start of behavioral testing. Tetrabenazine (TBZ) [(*R,R*)-3-Isobutyl-9,10-dimethoxy-1,3,4,6,7,11b-hexahydropyrido[2,1-*a*]isoquinolin-2-one] (Sigma-Aldrich), was dissolved in a 5% (v/v) dimethylsulfoxide (DMSO) solution mixed with saline and pH adjusted with 1 N HCl to bring the final solution to pH 6.5. TBZ was administered 100 min before testing. Citalopram, fluoxetine, SB 242084 (Tocris Bioscience, Bristol, UK) were dissolved in 0.9% (w/v) saline at the required concentrations and administered IP 30-min before the start of behavioral testing. All injections were administered at an injection volume of 10 ml/kg and treatment patterns were specified according to a Latin square.

Cohort 1 (N=15) was tested with citalopram, TBZ (1.5 and 3.0 mg/kg) and bupropion (5.0 and 10.0 mg/kg). Cohort 2 (N=15) was tested with SB204048 (0.25 and 0.50 mg/kg) and fluoxetine. Cohort 3 (N=15) was tested with TBZ (6 mg/kg) and SB204048 (0.5 and 1.0 mg/kg). A minimum of 3 drug-free days (5 days for TBZ) were required following completion of each Latin square. Animals were excluded if less than 50% of responses were made to the S+ for all the doses within a test. Thus, one animal was excluded from the TBZ (6 mg/kg) test, and one animal was excluded from the SB204048 (0.25 and 0.50) test. One saline 0.9% (w/v) injection was administered one week before the first pharmacological treatment cycle to habituate animals to handling and injection.

### Apparatus

All training and testing was performed in standard Bussey-Saksida mouse touchscreen chambers (Campden Instruments Ltd, Loughborough, UK). These chambers have been described in detail elsewhere (Horner et al. 2013; Heath et al. 2015). Briefly, the trapezoidal operant arena is housed inside a sound-attenuating chamber with a touchscreen mounted at one end and a reward delivery magazine at the other. A speaker and house light are mounted above the arena with a video camera for behavioural monitoring. Behavioral schedules are programmed using ABET II software (Campden Instruments Ltd, Loughborough, UK) with details provided in the subsequent sections.

### Habituation and initial operant training

On the day before habituation to the touchscreen chambers, animals were exposed to the liquid reward (strawberry milkshake (Yazoo, FrieslandCampina UK Ltd, Horsham, UK)) in their home cages to reduce neophobia by placing small bowls containing the milkshake in the cages overnight. On the day following the milkshake pre-exposure, animals were habituated to the chambers for 20 minutes. 200 µl of milkshake was provided in the reward collection magazine at the start of the habituation session with no programmed consequences.

Following habituation to the chambers, animals were trained to touch the screen. This procedure was carried out as per the standard ‘initial touch’ schedule (Horner et al. 2013; Heath et al. 2015). At the beginning of the session, one of two screen locations was illuminated and remained so until 30 seconds had elapsed, at which point the screen stimulus was removed, a tone (1000 ms, 3 kHz) was issued, the magazine was illuminated and 20 μL of liquid reward delivered (feeder pulse: 800 ms). After reward collection, the magazine light was turned off and a 5-s inter-trial interval (ITI) followed. A new trial then began. If the mouse touched the stimulus location while illuminated, the stimulus was immediately turned off, the tone issued, and the magazine illuminated. A triple delivery of reward was provided on these trials. Animals were considered successfully trained on this phase once 30 rewards were collected during a session.

### CJB task

The CJB task consisted of four stages: stages 1-3 provided incremental training to touch and discriminate between the rewarded (S+) and non-rewarded (S−) screen stimuli, and stage 4 provided the CJB assessment.

#### Stage 1: Learning to touch

In this stage, mice were trained to touch a white square presented within a white-outlined frame centred on the screen, with the bottom of the stimulus positioned 2 cm above the grid floor. The stimulus presentation duration (SD) in this stage was 10s, with an additional 0.5s limited hold (LH) period immediately following stimulus offset during which a response within the white frame would be considered valid. The LH period was instantiated in the schedule to enable capture of otherwise successful responses initiated immediately before stimulus offset (Kim et al. 2015).

If a successful touch response was made within the white frame during the SD or LH period, the white square stimulus was removed from the display (if present) and 20 µl of strawberry milkshake was delivered, along with a 1000 ms tone and illumination of the magazine light. A head entry into the magazine to collect the reward turned off the magazine light and initiated an inter-trial interval (ITI) of 2 s. A touch within the white frame during the ITI period (‘blank touch’) reset the ITI, so delaying the start of the next stimulus presentation. The criterion for stage 1 was for mice to earn 60 rewards within a 45 min session. The session was terminated either when 45 min had elapsed or when the maximum number of rewards (60) had been collected. Upon successful attainment of the criterion, animals progressed to Stage 2.

#### Stage 2: Positive Stimulus (S+) introduction

In stage 2, the white square stimulus was replaced by an S+ stimulus and the SD was reduced to 4 s (with a LH of 0.5 ms, thus animals had 4.5 ms to respond) an SD characterized by a low attentional load (Kim et al. 2015) to reduce as far as possible the effects of drugs on attention being the driver for any observed behavioural output. The S+ stimulus was either a horizontal or vertical line stimulus and was counterbalanced across individuals. From stage 2 onwards, a head entry into the magazine for reward collection initiated a brief ingestion delay (ID) period of 4 s before proceeding to the next ITI. The session was terminated either when 45 min had elapsed or when the maximum number of rewards (80) had been collected All other parameters and performance criteria were identical to those of stage 1.

#### Stage 3: Negative stimulus (S−) introduction and stimuli discrimination training (DT)

In stage 3, the S− stimulus was introduced and randomly presented in 50 % of trials. The ITI was also increased to 5 s. A touch to an S− stimulus during the SD or LH periods resulted in stimulus removal (if present) followed by an ITI period prior to a correction trial. On correction trials, the presented stimulus was always the S− and consecutive correction trials continued until no response to an S− was made during the SD or LH periods. Correction trials, together with ‘blank touches’ resetting the ITI (see stage 1), were included to discourage non-selective responding to the stimulus and screen, respectively. A response to the S− was followed by a correction trial, along with a 1000 ms tone and a 1000 ms flashing light. All other settings were identical to those of stage 2. All animals completed a minimum of seven sessions in stage 3 to fifteen an acquisition curve.

The proportions of responses and mistakes were calculated for the discrimination training as follows:

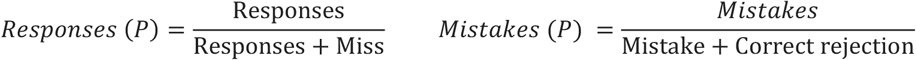

A response was recorded when a touch to the rewarded stimulus S+ was made and a miss was recorded when no response was made to the S+. A mistake was recorded when a response was made to the S− and a correct rejection was recorded when animals withheld responding to the S− (See **Figure 1**). The performance criteria for this stage required animals to achieve ≥75% responses. and ≤ 25% mistakes to yield a d’ ≥1.34 (see Data Analysis section).

**Figure 1.**
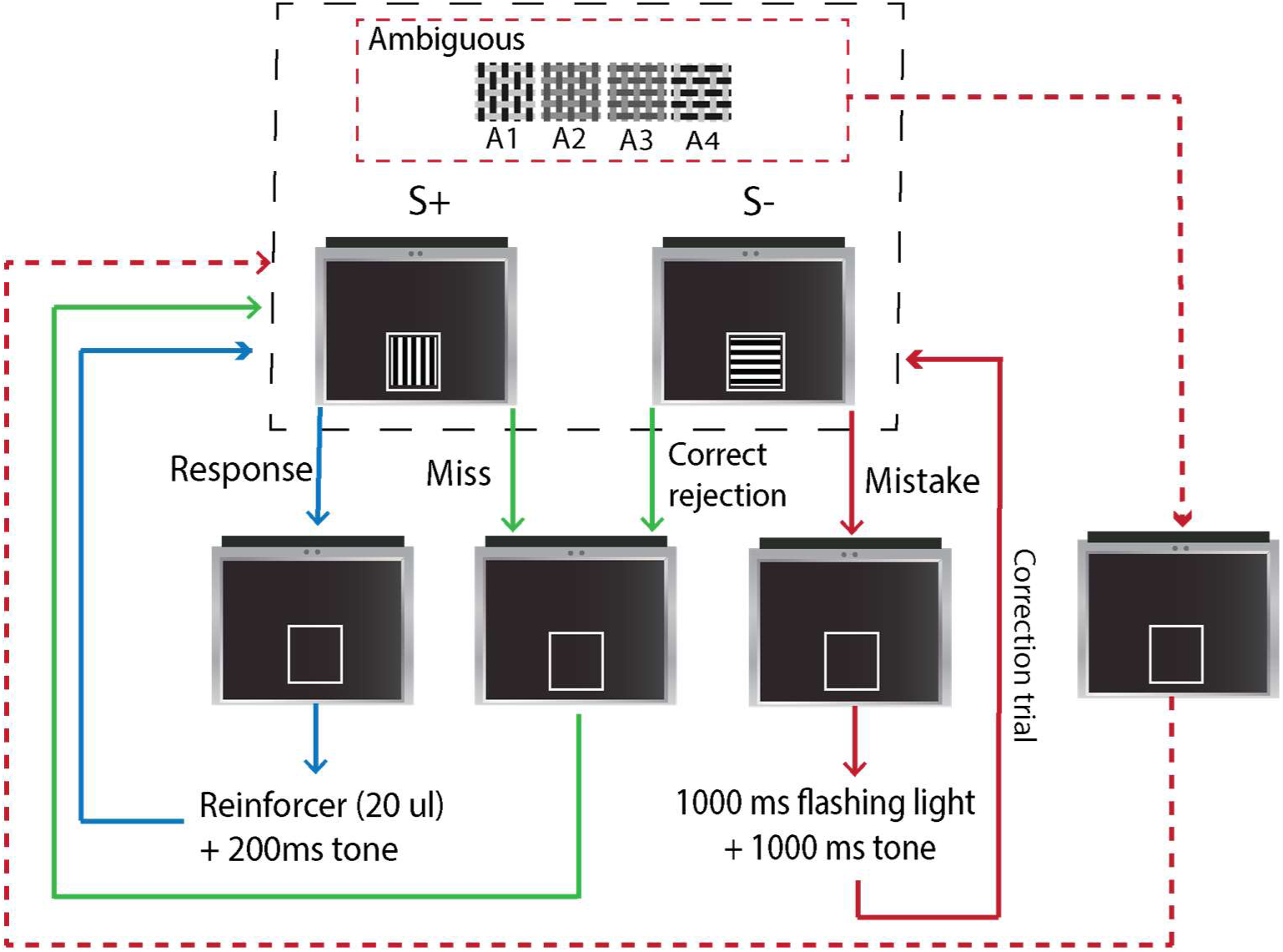
Discrimination training (Stage 3): On each trial, the S+ stimulus associated with reward and 200 ms tone delivery or the S− associated with 1000 ms flashing house light and tone delivery was presented within a white frame in the centre of the screen: After touching the stimulus a blank screen with a central white frame was presented during an inter-trial interval (ITI) preceding the next trial. Blue arrows indicate a touch to the S+ (Response). Green arrows indicate no response to the S+ (Miss) or no response to the S− (Correct rejection). Red arrows indicate a response to the S− (Mistake). A Response to the S+ resulted in reward delivery, with reward collection triggering a brief (4 s) ingestion delay period prior to the ITI. A miss or a correct rejection was followed by a new trial ITI, while a mistake was followed by a 1000 ms flashing house light, a tone and a correction trial. **Ambiguous stimuli introduction: Cognitive Affective Bias (CJB) assessment:** Ambiguous stimuli were randomly intermixed with the S+ and S− with no programmed consequences. The consequences for S+ and S− responding remained the same as in the discrimination training stage (stage 3).

#### Stage 4: Ambiguous stimuli introduction and CJB assessment

CJB assessment consisted of a discrimination task session as described in stage 3, with the addition of four ambiguous stimuli with intermediate features between the S+ and the S−.

The ambiguous stimuli were named A1 (most similar to S+), A2, A3 and A4 (most similar to S−). Within a test session, 40 ambiguous stimuli (10 of each) (see Figure 1) were intermixed with 50 S+ and 50 S− presentations. The proportion of responses and latencies to respond to each stimulus were recorded as the main measures as in Bethel et al., 2012. The other variables recorded during the sessions are specified in the Data Analysis section.

Animals were pre-exposed to CJB for one session before starting with pharmacological testing to ensure the pattern of responses to ambiguous stimuli was congruent with the level of similarity with the S+ or S-, as observed in other ambiguous interpretation-based CJB tasks (Harding et al. 2004; Enkel et al. 2010; Bethell et al. 2012). After this first CJB exposure, animals were re-baselined on DT (stage 3) and then tested on CJB after drug administration following a within-subjects design, leaving 3-5 days washout between tests while being re-baselined on DT (**Figure 2**).

**Figure 2.**
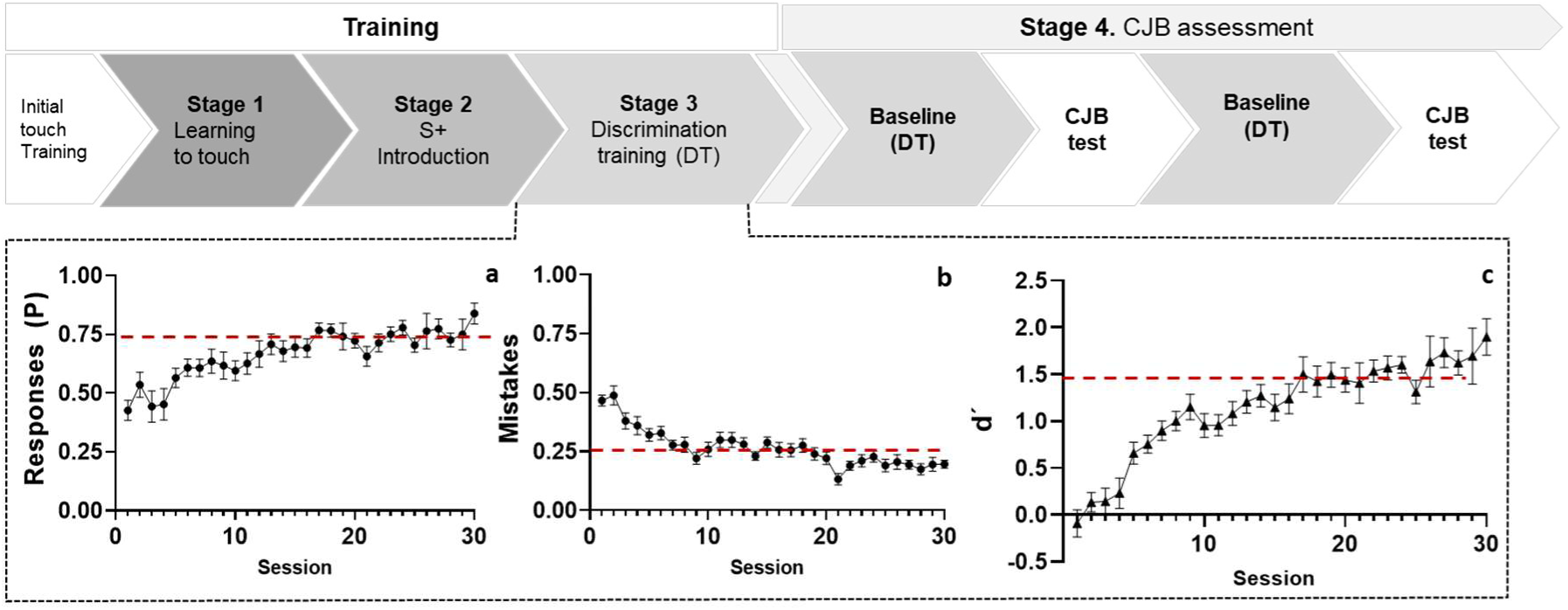
Experimental timeline. Initial-touch training was followed by pre-CJB training (Stage 1 to Stage 3). After reaching the criteria in Stage 3 (Discrimination training (DT)), animals were exposed to the CJB schedule (*) to test ambiguous stimuli feasibility to elicit an ambiguous-dependent pattern of responses and to reduce task novelty before pharmacological screening. After this, animals were re-baselined with the DT schedule and tested in CJB after the administration of drugs. In between tests, animals were re-trained in DT to maintain a stable baseline. Figures **a**, **b,** and **c**, represent the proportion of responses to S+, the proportion of mistakes, and the d’ value during Stage 3 (DT) for a representative group of animals (Cohort 3).

### Data analysis

For response performance profiling, the proportions of responses (Responses (P)) and Mistakes (P) made by each animal were calculated for each of the control trials (S+ and S–), and the ambiguous stimuli trials (A1, A2, A3, A4) delivered within the behavioural session. However, relying exclusively on these two parameters may lead to misinterpretation of task performance. For example, a high proportion of responses does not always indicate a good performance profile, indeed, when combined with an equally high mistake proportion, it is likely indicative of high levels of non-selective responding. For this reason, and to ensure that the proportions of hits to ambiguous stimuli were not biased by the effects of the administered drugs on the ability of the animals to discriminate or withhold responses, composite measures of Responses and Mistakes were used based on signal detection theory, yielding an index of sensitivity (d’) and an index of response bias (c). Sensitivity refers to the perceptual discriminability between the S+ and S−; i.e., higher values indicate better visual discrimination. Response bias refers to the criterion or willingness to make responses, e.g. conservative (high c) or liberal (low c) (Schulz et al. 2007; Kim et al. 2015).

The formulae for the calculation of these parameters are as follows:

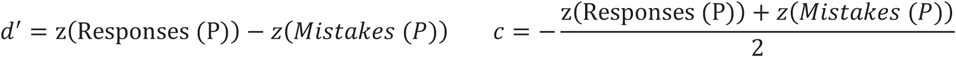

Prior to performing these calculations, all responses with a latency of 100 ms or less were removed from the raw data as they were likely to reflect unspecific responses. Additionally, the following measures were recorded: ‘blank’ touches to the empty frame during the ITI and proportion of correction trials.

Responses (P) were analyzed by two-way repeated measures ANOVA, followed by Sidak’s post hoc for multiple comparisons. Latencies to respond were analyzed using mixed effects ANOVA to account for missing values (withheld responses). d’, c, correction trials, and blank touches were analyzed with a paired samples t-test when only two doses were compared and by repeated measures ANOVAs when analyzing and comparing the effect of three or more doses.

Analyses and graphs were carried out with GraphPad Prism 8.4.3 (GraphPad Software, San Diego, California USA).

## Results

### Bupropion increased responses to ambiguous stimuli without affecting sensitivity and discrimination indexes

To investigate whether bupropion increased responses to ambiguous stimuli compared to vehicle treatment, an independent repeated-measures ANOVA was conducted and it revealed significant main effects of bupropion treatment and stimulus type on the proportion of responses [F (2, 42) = 5.068, p=0.0433; F (3.743, 157.2) = 190, p<0.0001, respectively], as well as a significant stimulus type x treatment interaction [F (10, 210) = 1.925, p=0.0433] (**Figure 3 a**). Post hocs revealed a significant increase in the proportion of responses to ambiguous stimulus A3 after 10 mg/kg bupropion (p=0.0017) and a significant increase in the proportion of responses after 5 mg/kg bupropion to ambiguous stimulus A4 (p=0.0324) compared with vehicle treatment. There were no differences in the proportion of responses to the S+ and S− between bupropion and vehicle treatment, thus the bupropion effect was specific under conditions of stimulus ambiguity.

**Figure 3.**
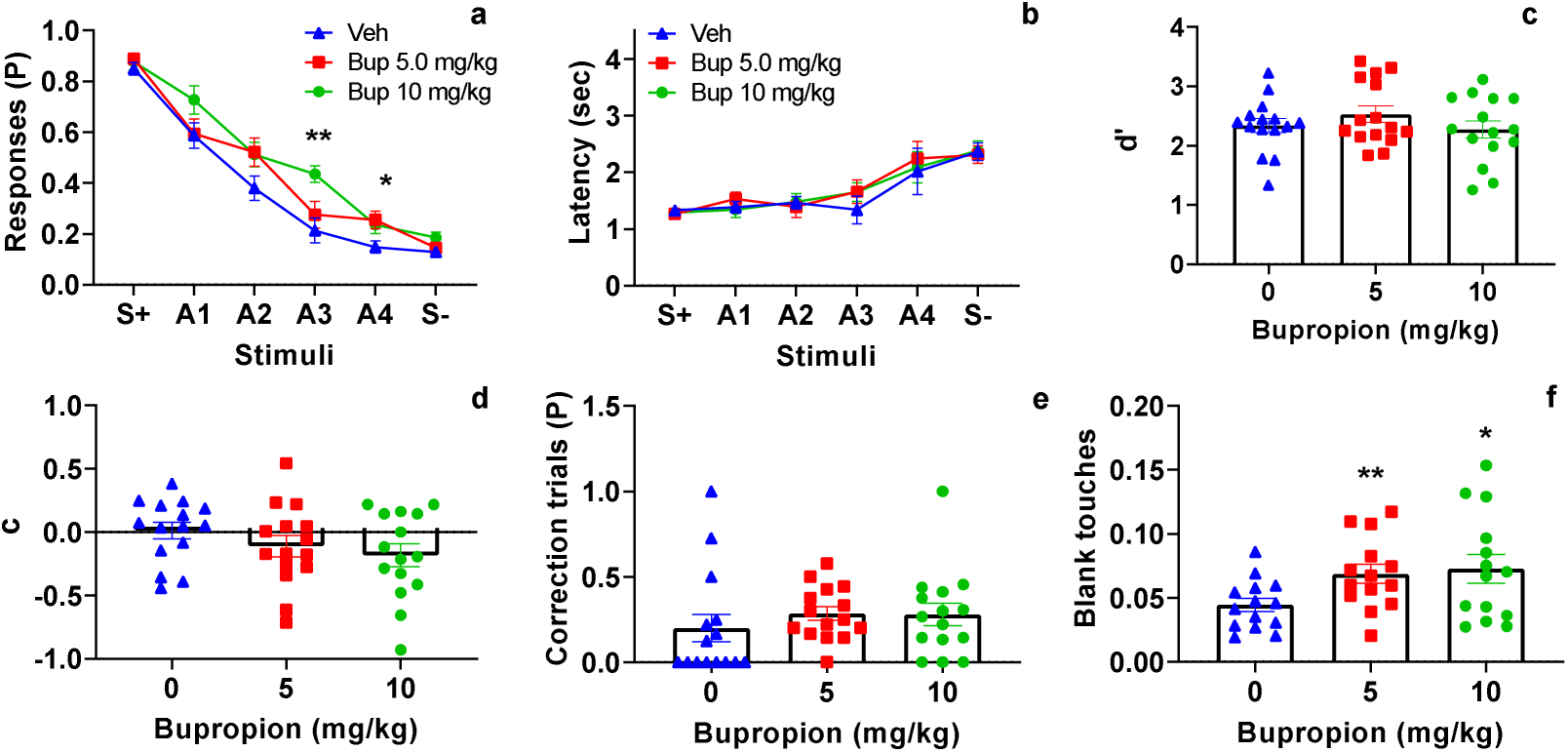
Effects of bupropion in the CJB task. Proportion of responses (**a**), latency to respond (**b**), d’ (**c**), c (**d**), the proportion of correction trials (**e**) and blank touches (**f**). *Data shown as mean (±SEM). **p<0.01, *p<0.05 significant differences from vehicle treatment*

Mixed effect analysis on latencies to respond did not show any effects of bupropion on this variable [F (2, 42) = 0.2316, 0.7943] nor a significant treatment x stimuli interaction [F (10, 202) = 0,2919, p=0.9825]. There was also a considerable effect of stimuli on latencies to respond [F(3.873, 49.96) = 0.2860, p<0.0001] (**Figure 3 b**).

Further repeated measures ANOVA analyses did not reveal effects of bupropion on the discrimination index (d’) [F (1,.940, 27.16) = 1.745, p=0.1943], or on the sensitivity index (c) [F (13677, 23.48) = 2,236, p=0.1358] (**Figure 3 c-d**). Although bupropion had a significant effect on blank touches [F (1.617, 21.02) = 6.867, p=0.0076] and both the dose of 5 and 10 mg/kg significantly increased these non-specific touches compared with vehicle (p=0.0038 and p=0.024, respectively) (**Figure 3 f**), it did not affect the proportion of correction trials [F (1.808, 25.31) = 0.7172, p=0.4844] (**Figure 3 e**), suggesting that although blank touches could be indicative of the stimulant effect of bupropion, this did not affect target-specific touches as shown by the lack of effect on the discriminatory and sensitivity indexes (d’ and c) (**Figure 3 c-d**). and the proportion of correction trials. (**Figure 3 e-f**).

### TBZ decreased ambiguous stimulus responses

Repeated measures ANOVA revealed significant effects of stimulus type [F (3.811, 99.07) = 84.56, p<0.0001] and treatment [F (1, 26) = 15.54, p=0.0005] and a stimulus type x treatment interaction [F (5, 130) = 2.808, p=0.0192] (**Figure 4 a**) on the proportion of responses under TBZ (6.0mg/kg). Post hoc analysis indicated a significant reduction in the proportion of responses recorded under TBZ for ambiguous stimuli A1 (p=0.0400) and A2 (p=0.0023) compared with vehicle treatment, with no effects of TBZ administration on the proportion of responses for the S+ or S− stimuli (p=0.2699 and, p=0.3172 respectively), indicating a pessimistic bias-like change in ambiguous stimuli perception under the drug.

**Figure 4.**
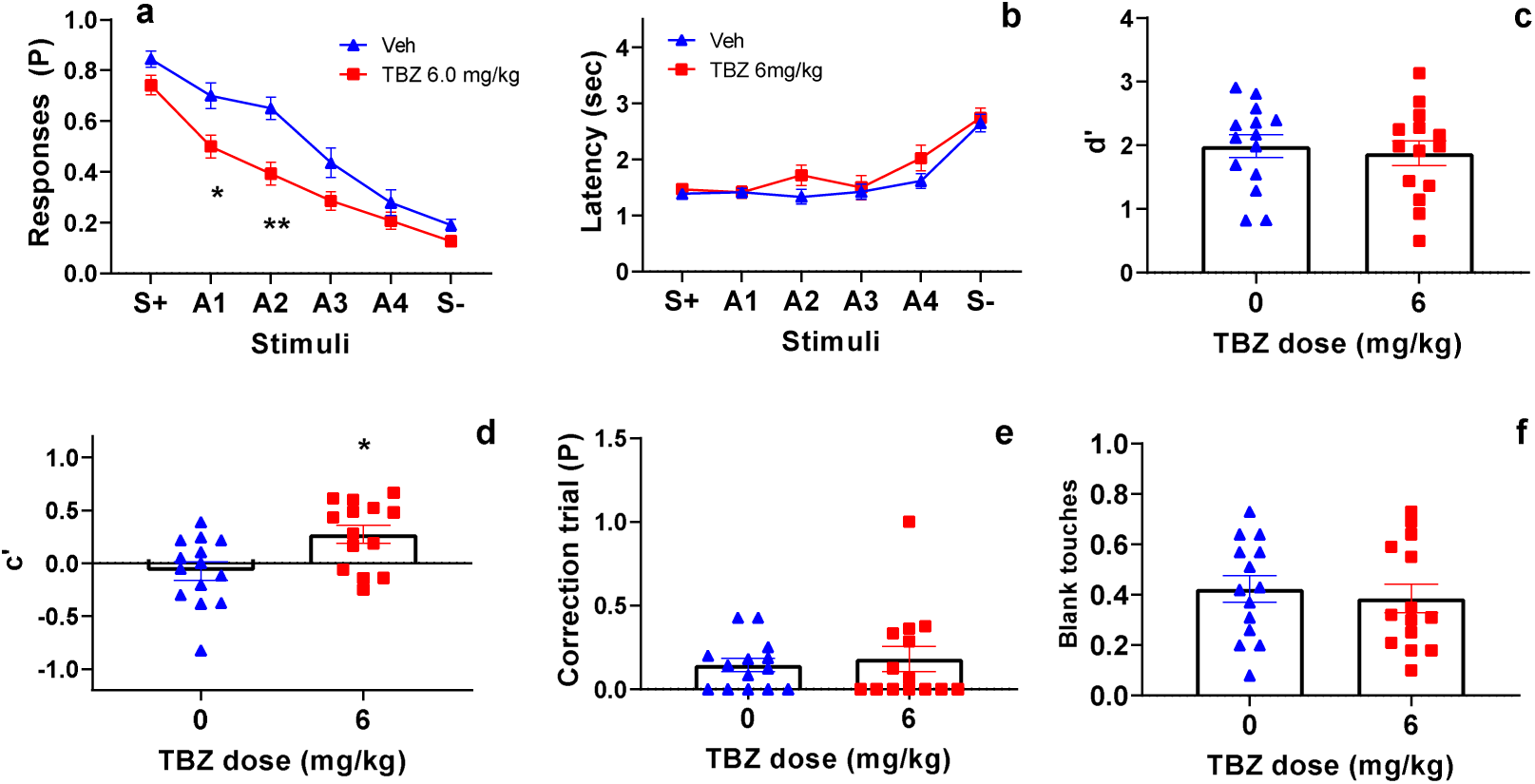
Effects of tetrabenazine (TBZ) in the CJB task. Proportion of responses (P) (**a**), latency to respond (**b**), d’ (**c**), c (**d**), proportion of correction of trials (**e**) and blank touches (**f**). *Data shown as mean (±SEM). **p<0.01 *p<0.05 significant differences from vehicle treatment*.

A significant main effect of stimulus type was detected on latency to respond [F (3,083, 46,24) = 10,27, p<0.0001], with no main effect of TBZ [F (2,503, 37.55) = 0.6048, p=0.5872] or a treatment x stimulus type interaction [F (5,978, 84,89) = 1,138, p=0.3477] detected for this variable (**Figure 4 b**). TBZ did not have effects on the discrimination index d’ (t(13)=0.8579, p=0.4066) but a significant effect on the sensitivity index (c) was detected (t(13)=1.446, p=0.0387) (**Figure 4 c-d**).

For the additional task performance measures, TBZ at this dose (6 mg/kg) did not have significant effects on the proportion of correction trials [t(13)=0.5902, p=0.5902] or blank touches [t(12)=0.8923 p=0.3885] (**Figure 4 e-f**).

### The SSRIs citalopram and fluoxetine and the 5HT-2C receptor antagonist SB201048 did not have effects on the CJB task performance

Two-way ANOVA indicated a main effect of stimulus type on the proportion of Responses observed under citalopram (F (4.160, 249.6) = 241,7, p<0.0001) but no main effects of treatment (citalopram: F (3, 60) = 0.7350, p=0.5352) or a treatment x stimulus type interaction (F (15, 300) = 0.2872, p=0.996) were observed (**Figure 5a**). With respect to latency to respond, a main effects of stimulus type was detected (F(3.083, 46.24) =10.27, p<0.0001) in the absence of a main effect of treatment (F (5.978, 84.89) = 1.138, p=0.3477) and a treatment x stimulus type interaction (F (5.978, 84.89) = 1.138, p=0.3986) (**Figure 5b**).

**Figure 5.**
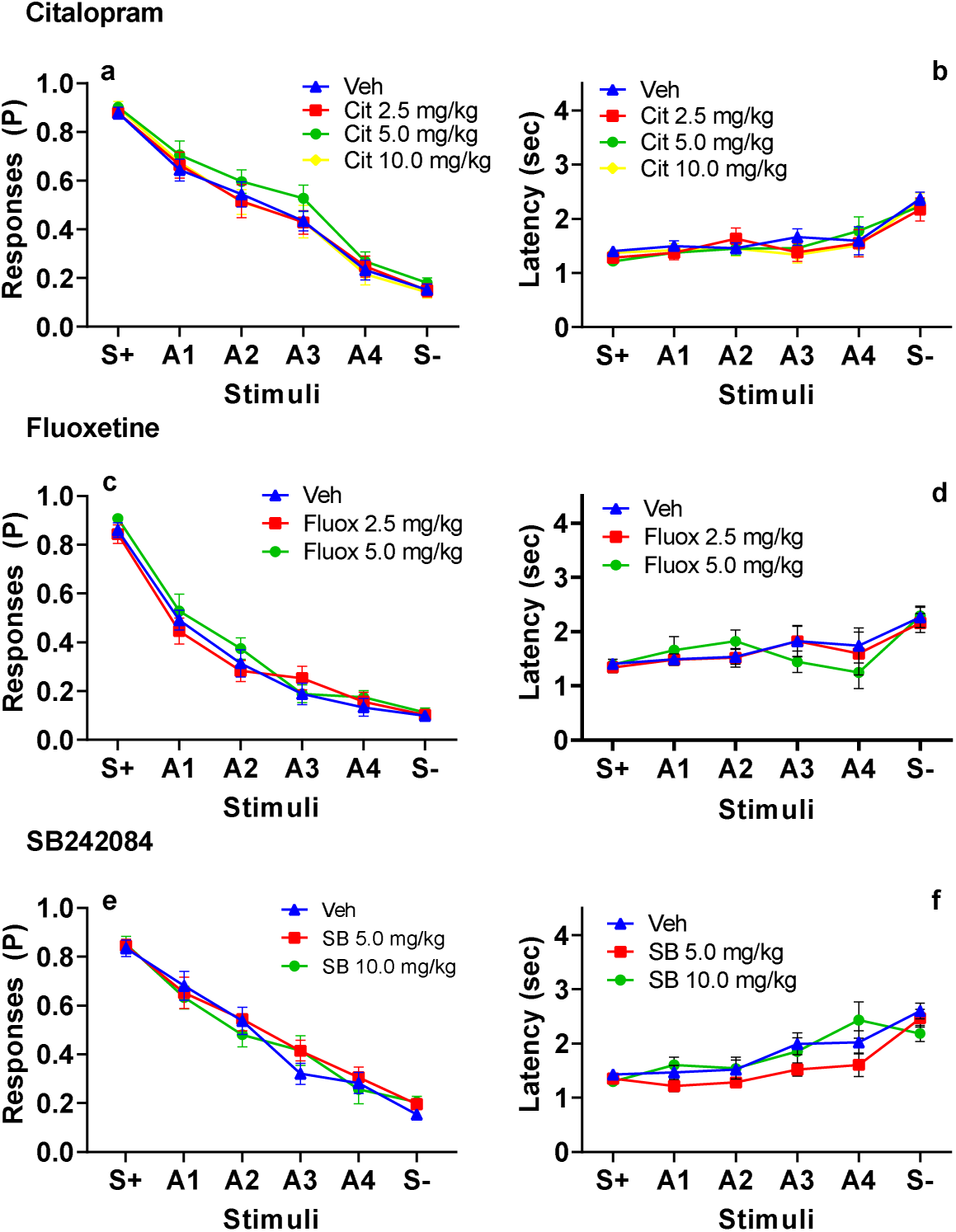
Effects of Citalopram, Fluoxetine and SB242084 on CJB task primary outcome measures. Proportion (P) of Responses (**a-c**) and Latency to respond (**d-f**) *Data shown as mean* (*± SEM*).

Similarly, while a main effect of stimulus type on the proportion of Responses to ambiguous stimuli was detected under fluoxetine (F (3.426, 154.2) = 227.5, p<0.0001), no main effect of fluoxetine on the proportion of responses to ambiguous stimuli (F (2, 45) = 0.5071, p=0.6056), or a treatment x stimulus type interaction (F (10, 225) = 0.8584) was detected(**Figure 5c**). With respect to latency to respond, a main effect of stimulus type was detected (F (2.772, 41.58) = 9.629, p<0.0001) in the absence of a main effect of treatment (F (2.467, 37.01) = 0.3824, p=0.7277) and a treatment x stimulus type interaction (F (4.794, 63.28) = 0.6581, p=0.6502) (**Figure 5 d**).

For the 5HT-2C receptor antagonist SB242084, two-way ANOVA indicated a main effect of stimulus type on the proportion of Reponses (F (3.678, 154.5) = 113.3, p<0.0001, **Figure 5e**) but no main effect of treatment (F (2, 42) = 0.156, p=0.856), or a treatment x stimulus type interaction (F(10, 210) = 0.655, p=0.765). The same pattern was observed on latency to respond after SB204048 did not have a main effect of treatment (F(1.186, 16.61)= 3.553, p=0.0714) and although the main effect of stimulus type was significant (F (2.680, 37.52) = 17.20, p<0.0001) the treatment x stimulus type interaction was not (F (4.89, 64.62) = 1.728, p=0.142) (**Figure 5f**).

Further CJB performance screening using a lower dose of SB204048 (0.25 mg/kg) did not detect any significant effect of treatment on any of the recorded variables (Supplementary Materials, Figure 2S and Table 2S).

For the additional task performance measures, both citalopram and fluoxetine did not have significant effects on the sensitivity index (d’), discrimination index (c), blank touches or the proportion of correction trials (See **Table 1**). SB242084 had effects only on the proportion of blank touches and Sidak’s *post hoc* showed that animals administered SB204048 at 1.0 mg/kg made more blank touches than when administered vehicle treatment (See **Table 1**).

**Table 1.** Effects of citalopram, fluoxetine, and SB204048 on d’, c, blank touches, and proportion of correction trials. *p<0.05, **p<0.01 significantly different from vehicle treatment.

| Drug/Dose<br>(mg/kg) | d' | c | Blank touches | Correction<br>trials (P) |
| --- | --- | --- | --- | --- |
| **Citalopram** |  |  |  |  |
| **0** | 2.45±0.12 | -0.10±0.11 | 0.05±0.005 | 0.19±0.05 |
| **2.5** | 2.54±0.17 | -0.03±0.12 | 0.07±0.011 | 0.29±0.08 |
| **5** | 2.36±0.12 | -0.21±0.09 | 0.07±0.01 | 0.24±0.05 |
| **10** | 2.71±0.25 | 0.06±0.11 | 0.06±0.01 | 0.24±0.05 |
| **F, p** | F (2.77, 41.55) =<br>0.75, p=0.52 | F (2.59, 38.91)=<br>0.57, p=0.62 | F (2.38, 35.74)=<br>2.84, p=0.06 | F (2.38, 35.73)= 0.57,<br>p=0.60 |
| **Fluoxetine** |  |  |  |  |
| **0** | 2.58±0.17 | 0.05±0.09 | 0.06±0.01 | 0.21±0.06 |
| **2.5** | 2.64±0.15 | 0.13±0.15 | 0.04±0.005 | 0.27±0.08 |
| **5.0** | 2.77±0.16 | -0.07±0.16 | 0.05±0.01 | 0.12±0.04 |
| **F, p** | F (1.96, 29.47) =<br>0.46,<br>p=0.63 | F (1.23, 18.43)=<br>1.22<br>, p=0.30) | F (1.73, 26.00)=<br>2.65, p=0.10 | F (1.41, 21.14)= 1.81,<br>p=0.19 |
| **SB242084** |  |  |  |  |
| **0** | 2.126±0.14 | 0.006±0.08 | 0.05±0.01 | 0.19±0.04 |
| **0.5** | 2.145±0.23 | -0.094±0.07 | **0.07±0.01\*** | 0.25±0.07 |
| **1.0** | 2.109±0.17 | -0.14±0.09 | **0.07±0.01\*\*** | 0.36±0.08 |
| **F, p** | F (1.720, 24.08)=0.03<br>p=0.96 | F (1.643,<br>23.00) = 2.259)<br>= 0.13 p=0.131 | **F (1.726, 24.16)=**<br>**6.37, p=0.008** | F (1.967, 27.53) =<br>3.253, p=0.054 |

## Discussion

In the present study, we developed and validated a Go/No Go CJB task for pro-depressant and antidepressant manipulation screening in mice using touchscreen technology that facilitates assessment across species (Nithianantharajah et al. 2015; Bethell 2016; Heath et al. 2019; Sullivan et al. 2021) thus enhancing the significant translational potential of the study of cognitive-affective bias (Robinson 2018); (Anderson et al. 2012).

Although there are multiple tasks available for assessing CJB in rats (Rygula et al. 2013, 2015; Papciak et al. 2013; Hales et al. 2014; Stuart et al. 2017; Drozd et al. 2019), the number of tasks available for assessing this psychological construct in mice (Novak et al. 2015) and particularly those based on operant schedules and featuring interpretation of ambiguous visual stimuli, is more limited (Krakenberg et al. 2019, 2020). The current study adapted a Go/No Go task used previously for assessing CJB via visual stimulus interrogation in non-human primates (Bethell et al. 2012).

The use of multiple ambiguous stimuli presented in an invariant spatial location within session in this CJB task ensures that observed responses to ambiguity are not biased by stimulus novelty, position or chance. This multi-stimulus design elicited a pattern of responding congruent with the level of ambiguity (i.e., similarity to S+ or S−) as shown in all the vehicle groups of the present study (**Figure 3-6 a**). Mice responded approximately 80-85% of the time to the S+ and approximately 15-20% to the S−. The proportion of responses to the ambiguous stimuli decreased as their similarity to the S+ decreased. This pattern of responding was almost identical to the touchscreen-based Go/No Go CJB task used in non-human primates (Bethell et al. 2012; Bethell and Koyama 2015) and to the first Go/No Go CJB task in rats using tone stimuli (Harding et al. 2004) which validates the feasibility of using visual stimuli and the non-human primate design in mice and emphasizes the potential for translation across species.

Latencies to respond to ambiguous stimuli were also as expected according to the literature in rodents, non-human primates, and humans (Harding et al. 2004; Bethell et al. 2012; Hales et al. 2022). As in previous Go/No go CJB task designs (Harding et al. 2004; Bethell et al. 2012), the latencies to respond to the S− were the longest compared with the other stimuli. Mice responded faster to the S+ and became slower as the stimuli presented increased in similarity to the S− and this pattern was identical across all the pharmacological studies. However, unlike the proportion of correct responses to each stimulus, the effect of affective-related manipulations on latencies to respond is more variable depending on the task design (Anderson et al. 2013; Barker et al. 2018).

The noradrenaline-dopamine reuptake inhibitor bupropion has antidepressant-like effects on classical models of depression like the FST (Carratalá-Ros et al. 2023), improves apathy in rodents (Randall et al. 2014) and also increases optimistic bias in humans (Walsh et al. 2018a, b). However, although selective noradrenaline uptake inhibitors like reboxetine, and serotonin/noradrenaline reuptake inhibitors like venlafaxine have been shown to increase optimistic bias in rats (Stuart et al. 2013), the effect of bupropion on cognitive-affective bias in rodents has not to our knowledge yet been studied. In the present study, this compound increased optimistic bias as shown by an increase in responses to ambiguous stimulus 3 (A3) at 10mg/kg and to ambiguous stimulus 4 (A4) at 5 mg/kg. However, bupropion had no effect on the percentage of responses to the S+ and S−., suggesting the effect of bupropion is specific to conditions of stimulus ambiguity (**Figure 3a**).

To further validate novel CJB task sensitivity to changes in mood, TBZ was utilised based on previous research showing pro-depressant effects of this drug on models of depression like the FST (Carratalá-Ros et al. 2023), on effort-related choice (López-Cruz et al. 2018; Yang et al. 2020)) and in cognitive-affective bias tasks (Stuart et al. 2017). In the present study, TBZ at 6 mg/kg induced a negative bias, manifested as a decrease in the proportion of responses to ambiguous stimuli without affecting the sensitivity index (d’). TBZ therefore did not affect animals’ sensitivity to the S+ and S− and so the observed responses were not driven by changes in how animals responded to the S+ or withheld responses to the S−. (**Figure 4 a, c**). TBZ did affect the response bias index parameter (c), making animals more conservative overall (**Figure 4 d**). The effect of TBZ resembles the pattern of results observed in similar Go/Go designs after pro-depressant manipulations like unpredictable housing in rats (Harding et al. 2004) and stressful routine vet checks in non-human primates in the primate version of our task (Bethell et al. 2012) as both manipulations reduced the proportion of responses to ambiguous stimuli.

TBZ did not significantly affect latencies to respond, although animals under TBZ were slightly slower in responding to ambiguous stimuli A2 and A4. This is congruent with the lack of effects of TBZ on this variable observed in other CJB tasks using high versus low reward as positive and negative discriminators (Hales et al. 2022).. Also, although previous studies using the primate version of this task showed that animals were slower at responding to ambiguous stimuli after stressful vet checks, the control group was environmentally enriched (Bethell et al. 2012); thus the effect of TBZ in this design may depend on subjects’ baseline latencies. Lastly, although high/moderate doses of TBZ could cause non-specific effects due to its effects on the monoamine system (Podurgiel et al. 2013; Yang et al. 2020) which could affect animals’ sensitivity to negative feedback after a mistake (i.e. correction trials) or decrease locomotor activity, at the dose used here, the drug did not affect the number of correction trials (**Figure 4e**) or decrease the number of blank touches (**Figure 4f**). Lower doses of TBZ (1.5 and 3.0 mg/kg) did not have significant effects on CJB task performance (Figure 1S and Table S1, Supplemental Materials).

The SSRIs citalopram and fluoxetine did not increase optimistic biases in the Go/No Go CJB test. This lack of effect is consistent with previous studies that suggest these antidepressants do not have effect on CJB tests when administered acutely in animals (Anderson et al. 2013; Golebiowska and Rygula 2017; Hales et al. 2017). This is also consistent with the effects of SSRIs in humans as they typically demonstrate therapeutic efficacy only weeks after treatment onset (Anderson et al. 2000) (although acute citalopram has been shown to increase happy face recognition in healthy volunteers (Murphy et al. 2009)). In rats, both acute fluoxetine and citalopram (1.0 mg/kg, IP) induced an optimistic bias in a bowl-digging task (Stuart et al. 2013). However, in operant Go/Go paradigms, low doses of citalopram (1 mg/kg) biased animals towards the positive interpretation of the ambiguous stimuli (tones) while high doses had the opposite effect (5-10 mg/kg) (Rygula et al. 2014). However, SSRIs have not previously been tested in Go/No-Go tasks involving ambiguous interpretation, making direct comparisons with the data presented here difficult.

SB242084, a 5-HT2C receptor agonist, has previously shown anxiolytic effects (Kantor et al. 2005), fast-onset antidepressant-like effects (Opal et al. 2014) and feedback sensitivity modulation in mice (Phillips et al. 2018). In the present study, the dose that modulated feedback sensitivity in our laboratory (1.0 mg/kg) (Phillips et al. 2018) did not affect ambiguous stimuli interpretation, nor the discrimination and sensitivity indexes. This dose (1.0 mg/kg) was observed to significantly increase blank touches which may reflect a non-specific stimulant effect (**Table 1**), which could explain the increasing trend in the proportion of correction trials. This trend aligns with earlier findings of reduced sensitivity to negative feedback following SB242084 administration (Phillips et al. 2018). The effect of SB242084 on latencies overall was not significant, however, a trend was observed with the dose 0.5 mg/kg after which animals responded to ambiguous stimuli A1-A4 (**Figure 5f**). Lower doses of SB242084 (0.25 and 0.5 mg/kg) did not have effects on CJB task performance (Figure S2 and Table S2, Supplemental Materials).

The present task may have certain limitations inherent to the Go/No-Go design. For instance, “pessimistic” withholding responses (i.e., “No-Go” responses) could potentially be influenced by factors such as motivational or locomotor activity. However, these concerns have been mitigated through the evaluation of discrimination and sensitivity indices and monitoring blank touches to control for non-specific effects on task performance. Future experiments should reduce the number of ambiguous stimuli to three to have a 50% ambiguous stimulus (equal feature overlap with the S+ and S−) as in CJB tasks used in other species which feature visual (Bethell et al. 2012) or auditory stimuli (Enkel et al. 2010; Rygula et al. 2014), which would also increase statistical strength.

The lack of effects of antidepressants could be also due to within-subject ‘optimism’ or ‘pessimism’ at baseline (Golebiowska and Rygula 2017). In the context of this task, this could explain why TBZ could induce the expected pro-depressant effect by decreasing responses to stimuli A1 and A2 (which elicited more responses in this task) and conversely why bupropion showed antidepressant effects on stimuli A3 and A4 which are the ambiguous stimuli that elicited fewer responses. Moreover, baseline responses in the TBZ experiment were higher for unknown reasons which may have facilitated the induction of pessimistic bias in this task. Future studies may incorporate reversal experiments in which antidepressant manipulations could be given to revert pessimistic biases induced by pro-depressant manipulations. Future studies should also include female mice as CJB baseline and intervention data derived solely from male animals severely limits translational value in the clinical context (Choleris et al. 2018; Warthen et al. 2020).

Despite these areas of improvement, the present design demonstrates that mice can respond in an ambiguity-dependent manner to visual stimuli as observed in other CJB tasks in other species. Unlike other tasks using the avoidance of a mild electrical shock, or a smaller reward as a negative outcome, the present task uses omission of reward as a negative outcome and still replicates the ambiguity-dependent pattern of responses observed in other tasks (Harding et al. 2004; Enkel et al. 2010) thus demonstrating that the contingencies, and the stimuli used were highly effective. This task therefore has been shown to work across species (Bethell et al. 2012).

Unlike other mice touchscreen CJB tasks (Krakenberg et al. 2019, 2020), the inclusion of stimuli exhibiting various levels of ambiguity presented in a fixed location corroborates the view that responses to the stimuli here are not biased by stimuli novelty, risk-taking behavior, spatial location or chance and, stimuli were also counterbalanced to avoid stimuli preference that could bias the responses. The present task was also shown to be sensitive to the effects of antidepressant and pro-depressant drugs, thus it establishes the basis for further optimization and screening of a wider range of pharmacological or non-pharmacological manipulations, including in the context of the assessment of animal welfare (Bethell et al. 2012; Lopez-Cruz et al. 2021).

Finally, the novel Go/No-Go CJB task for mice can be integrated with over 20 other touchscreen-based tests (Sullivan et al. 2021), expanding the range of tools available for assessing affective state-related constructs (Phillips et al. 2018; Krakenberg et al. 2019, 2020) in this apparatus. This facilitates the deployment of focused ‘suites’ of touchscreen assessments within single cohorts of animals, enabling within-subject assessment of several behavioral domains under the same conditions and using the same manipulanda. This, combined with the use of reward omission as a negative outcome (rather than, e.g., mild electrical shocks), strongly aligns with the 3Rs Principles of Reduction and Refinement (Lopez-Cruz et al. 2021).

## ACKNOWLEDGMENTS

This study was supported by an NC3Rs project grant (NC/N001451/1). We gratefully acknowledge Dr. Paul A. S. Sheppard for helpful comments on an earlier version of the manuscript and Esmin Unaran and Rachel Bedwin for their help in performing behavioural experiments.

## AUTHOR CONTRIBUTION

LLC designed and conducted the experiments and wrote the first draft of the manuscript. BUP supported with experiments. TJB, CJH, and LMS supervised the project. All authors reviewed, critiqued, and commented on drafts of the manuscript and gave final approval for the submission.

