## Supplementary Materials for "Characterisation of a touchscreen-based Go/No Go task for assessing cognitive judgement bias in mice: a new translational tool for affective state disorder drug screening"

**Tetrabenazine at low doses did not have effects on the CJB task performance.**

Two-way ANOVA indicated a main effect of stimulus type on the proportion of Responses observed under TBZ (F(3.437, 96.24) = 114.0, p<0.0001) but no main effects of treatment (F (1, 28) = 0.014, p=0.9152) or a treatment x stimulus type interaction (F(5, 140) = 1,348,p=0.2477) were observed (**Figure 1S a).** Concerning latency to respond, a main effect of stimulus type was detected (F (2.610, 69.43) = 11.19, p<0.0001) in the absence of a main effect of treatment (F (1, 28) = 1,835, p=0.1863) and a treatment x stimulus type interaction (F (5, 133) = 0,7301, p=0,6021) **(Figure 1S b**).

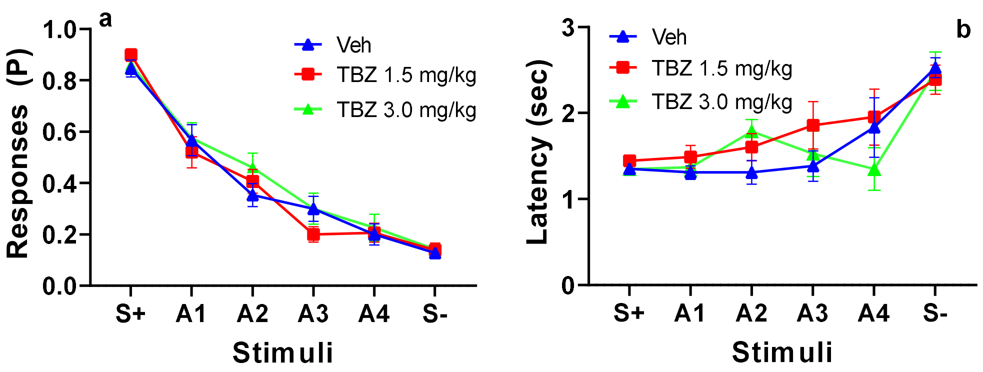


**Figure 1S. Effects of low doses of TBZ on CJB task primary outcome measures.** Proportion (P) of Responses (**a**) and Latency to respond (**b**) *Data shown as mean (± SEM).*

**Other variables:**

| **Drug/Dose (mg/kg)** | **d'** | **c** | **Blank touches** | **Correction trials (P)** |
|---|---|---|---|---|
| **T BZ** | | | | |
| **0** | 2.29±0.14 | 0.06±0.11 | 0.06±0.01 | 0.23±0.09 |
| **1.5** | 2.57±0.12 | -0.05±0.10 | 0.05±0.01 | 0.22±0.06 |
| **3.0** | 2.36±0.15 | 0.05±0.10 | 0.05±0.01 | 0.14±0.04 |
| **F, p** | F (1.77, 24.77) = 1.63, p=0.22 | F (1.64, 24.67)= 0.77, p=0.45 | F (1.74, 24.40) = 1.74 p=0.20 | F (1.36, 19.07) = 0.48, p=0.56 |

**Table 1S.** Effects of TBZ at low doses on d´, c, blank touches, and proportion of correction trials.

**SB at low doses did not have effects on the CJB task performance.**

Two-way ANOVA indicated a main effect of stimulus type on the proportion of Responses observed under SB (F(4.101, 172.2) = 115.4, p<0.0001) but no main effects of SB (F(2, 42) = 0.892, p=0.418) or a treatment x stimulus type interaction (F (10, 210) = 0.448, p=0.959) were observed (**Figure 2S a).** With respect to latency to respond, a main effects of stimulus type was detected (F (3.481, 48.73) =

18.10, p<0.0001) in the absence of a main effect of treatment (F (1.423, 19.93) = 0.711, p=0.4573) and a treatment x stimulus type interaction (F (4.311, 50.44)= 0.7946, p=0.5425) **(Figure 2S b**).

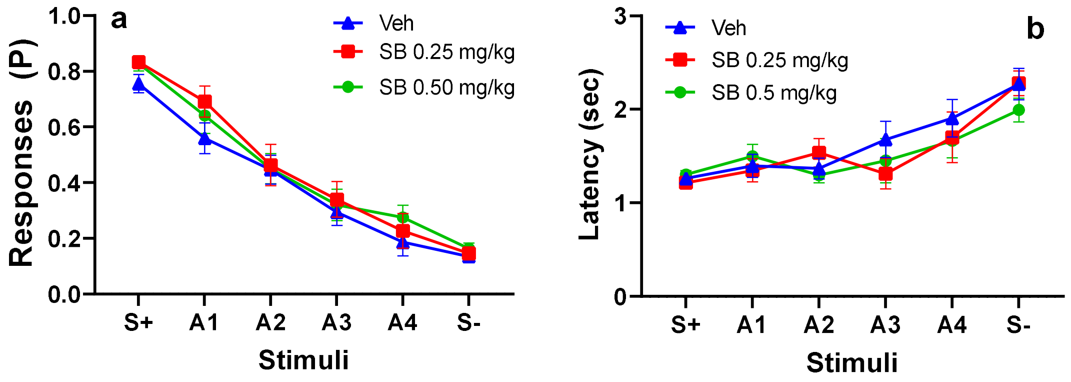


**Figure 2S. Effects of low doses of SB242084 on CJB task primary outcome measures.** Proportion (P) of Responses (**a**) and Latency to respond (**b**) *Data shown as mean (± SEM).*

***Other variables:***

| **Drug/Dose (mg/kg)** | **d'** | **c** | **Blank touches** | **Correction trials (P)** |
|---|---|---|---|---|
| **SB** | | | | |
| **0** | 1.96±0.143 | 0.21±0.09 | 0.05±0.01 | 0.20±0.05 |
| **0.25** | 2.07±0.11 | **0.02 ±0.09*** | 0.06±0.01 | 0.18±0.05 |
| **0.5** | 2.15±0.15 | 0.04±0.08 | 0.06±0.01 | 0.23±0.07 |
| **F, p** | F (1.87, 26.18) = 0.72, p=0.50 | F (1.823, 25.53) = 4.06, **p=0.03** | F (1.51, 21.14) = 3.22, p=0.07 | F (1.73, 24.32) = 0.19, p=0.79 |

**Table 2S.** Effects of citalopram, fluoxetine, and SB204048 on d´, c, blank touches, and proportion of correction trials. *p<0.05, significantly different from vehicle treatment.
